# A reappraisal of the default mode and frontoparietal networks in the common marmoset brain

**DOI:** 10.1101/2023.11.28.569119

**Authors:** Takuto Okuno, Noritaka Ichinohe, Alexander Woodward

## Abstract

In recent years the common marmoset homologue of the human default mode network (DMN) has been a hot topic of discussion in the marmoset research field. Previously, the posterior cingulate cortex regions (PGM, A19M) and posterior parietal cortex regions (LIP, MIP) were defined as the DMN, but some studies claim that these form the frontoparietal network (FPN). We restarted from a neuroanatomical point of view and identified two DMN candidates: Comp-A (which has been called both the DMN and FPN) and Comp-B. We performed GLM analysis on auditory task-fMRI and found Comp-B to be more appropriate as the DMN, and Comp-A as the FPN. Additionally, through fingerprint analysis, a DMN and FPN in the tasking human was closer to the resting common marmoset. The human DMN appears to have an advanced function that may be underdeveloped in the common marmoset brain.

## Introduction

The default mode network (DMN), a network of brain regions in humans, is activated when a person is at rest, during introspective moments like remembering the past, envisioning the future, or when considering the thoughts and perspectives of other people [1] [2]. This prominent network has also been observed in other animal species such as the chimpanzee [3], macaque [4] [5], common marmoset [6] [7] [8], rat [9], and mouse [10]. The DMN can be extracted through several neuroimaging techniques, such as independent component analysis (ICA) of resting-state (rs-) fMRI (functional magnetic resonance imaging) [11] [12], seed-based connectivity analysis (SCA) of rs-fMRI [12], or by task-induced deactivation of general linear model (GLM) analysis of task-fMRI [1] [13]. Usually, the ICA approach is favoured for the extraction of large brain network components in humans and non-human primates. The areas of the DMN are widely agreed upon for the human brain (see Table 1, Fig. 1, and for example, [1]). However, for the common marmoset (Callithrix jacchus), a non-human primate, the homologue of the human DMN has been a hot topic of discussion in recent years in the marmoset research field.

**Fig. 1.**
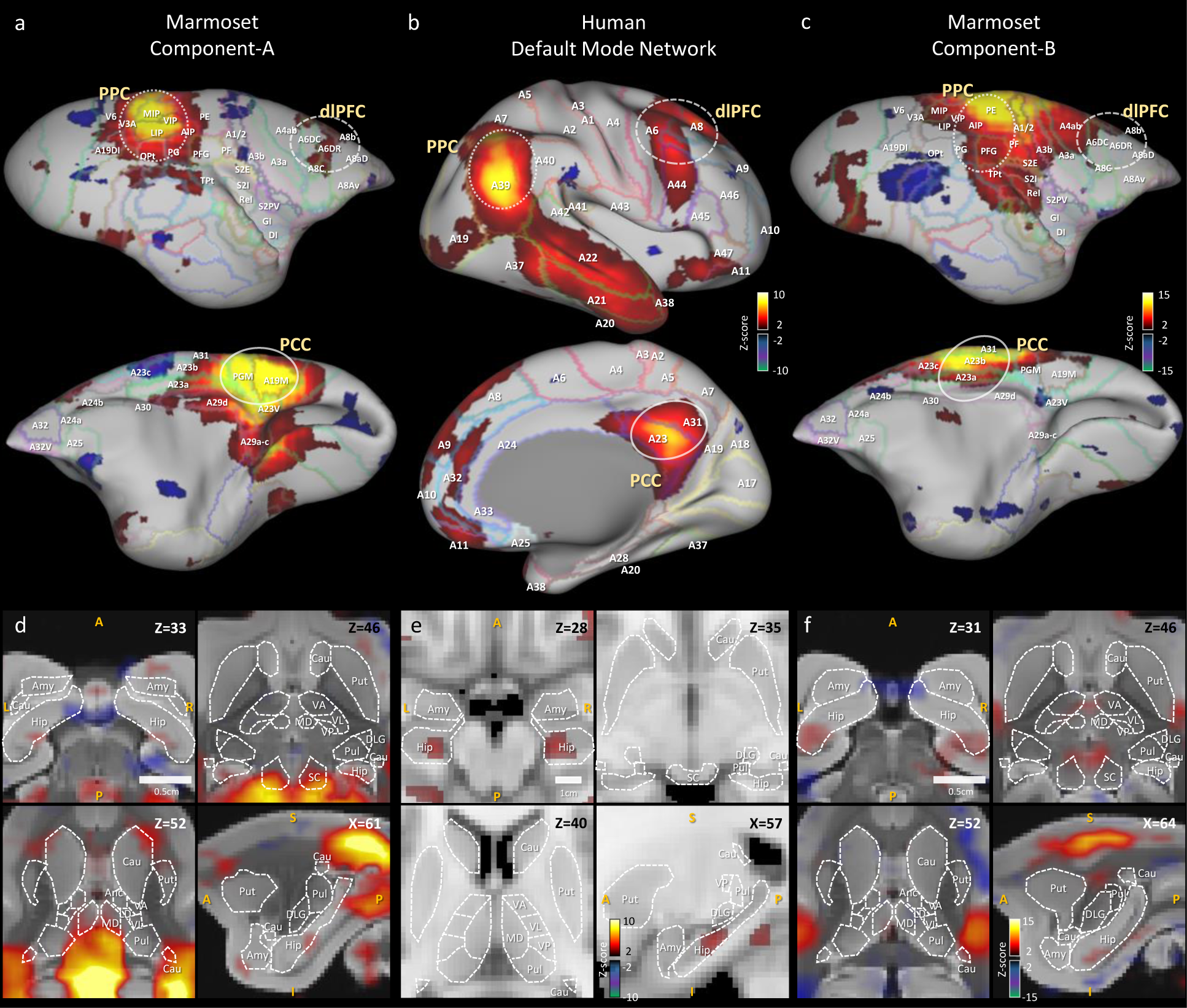
Human default mode network component and awake marmoset ICA components. **a**, Right cortical surface of the marmoset, (top) lateral side, (bottom) medial side. Awake resting-state marmoset ICA component-A, selected from 30 components, mapped onto the brain surface. Z-score range is 2 to 15 for positive, -2 to -15 for negative. **b**, Right cortical surface of the human brain. Human resting-state default mode network is mapped onto the surface. Z-score range is 2 to 10 for positive, - 2 to -10 for negative. **c**, Right cortical surface of the marmoset brain. Awake resting-state marmoset ICA component-B, mapped onto the brain surface. **d**, Horizontal views (top left, right and bottom left) and a sagittal view (bottom right) of awake marmoset ICA component-A. Scale bar shows 0.5cm. Z-score range is from 2 to 15. **e**, Horizontal views (top left, right and bottom left) and a sagittal view (bottom right) of human resting-state default mode network component. Z-score range is from 2 to 10. Scale bar shows 1cm. **f**, Horizontal views (top left, right and bottom left) and a sagittal view (bottom right) of awake marmoset ICA component-B. Abbreviations are Caudate (Cau); putamen (Put); hippocampus (Hip); amygdala (Amy); superior colliculus (SC); thalamus anterior nuclear complex (Anc), laterodorsal (LD), mediodorsal (MD), ventral anterior (VA), ventral lateral (VL), ventral posterior (VP), pulvinar (Pul) and lateral geniculate (DLG).

**Table 1.**
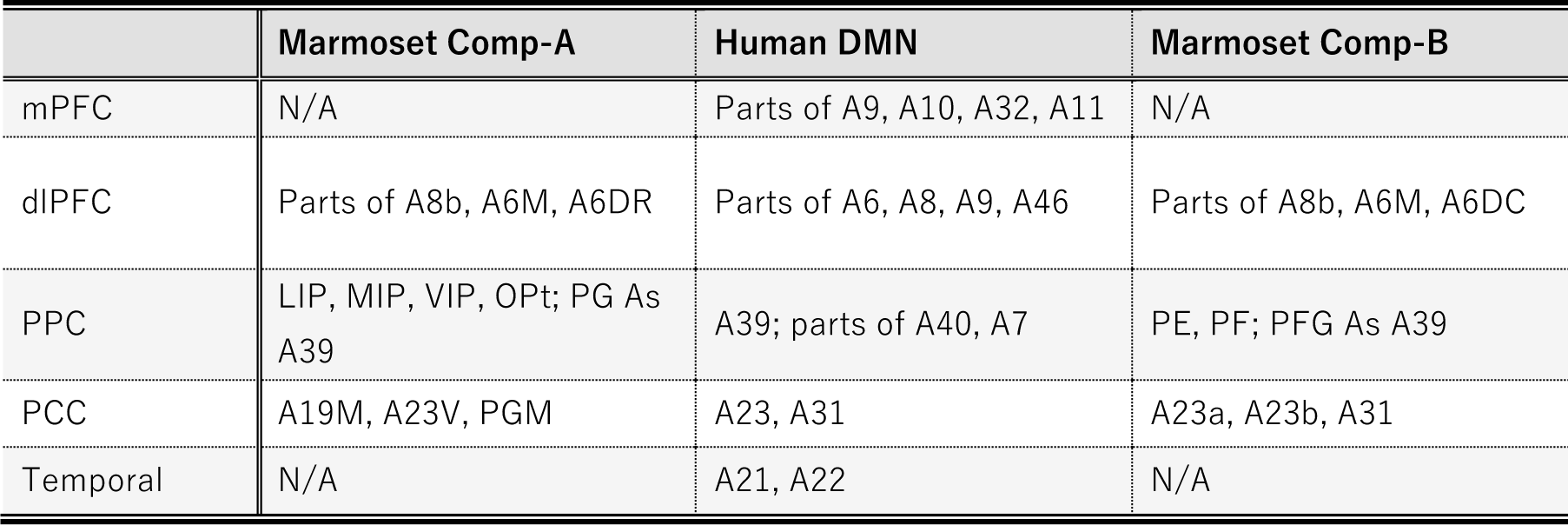
Default mode network regions under investigation and their areas.

|  | Marmoset Comp-A | Human DMN | Marmoset Comp-B |
| --- | --- | --- | --- |
| mPFC | N/A | Parts of A9, A10, A32, A11 | N/A |
| dIPFC | Parts of A8b, A6M, A6DR | Parts of A6, A8, A9, A46 | Parts of A8b, A6M, A6DC |
| PPC | LIP, MIP, VIP, OPt; PG As A39 | A39; parts of A40, A7 | PE, PF; PFG As A39 |
| PCC | A19M, A23V, PGM | A23, A31 | A23a, A23b, A31 |
| Temporal | N/A | A21, A22 | N/A |

The DMN of the common marmoset was first described by Belcher et al. [6]. Group-ICA was applied to rs-fMRI sessions, and was defined as consisting of the retro-splenial and posterior cingulate cortex (PCC) region (A23, A31, A29 and A30 areas), the dorsolateral prefrontal cortex (dlPFC) region (A6DR, A6DC and A8C areas), the posterior parietal cortex (PPC) region surrounding PE, PFG, PG, and the left intraparietal sulcus (LIP) and middle intraparietal sulcus (MIP). Ghahremani et al. [14] identified the same network component by group-ICA, but they instead defined it as the frontoparietal network (FPN). This was because it had previously been reported and identified as a frontoparietal network controlling saccades in resting-state network (RSN) studies of anesthetized macaques [15]. Liu et al. [7] refuted this and argued that this component is the DMN, because it was found that task-induced deactivation in visual-task fMRI occurs around the PCC (PGM and A19M areas) and PPC (LIP and MIP areas) regions. This definition was continued with in Tian et al. [16]. In later research, Hori et al. [8] applied fingerprint analysis [17] using several sub-cortical regions and found that this component was the closest to the DMN component obtained from human rs-fMRI, and therefore concluded it to be the DMN of the common marmoset. Ngo et al. [18] applied joint gradient analysis [19], and gradient 2 showed similarity between the resting human DMN and marmoset dlPFC-PCC-PPC network. Although these studies appear to have reached some consensus, some studies continue to use the FPN definition [20] [21]. Furthermore, there remains a large mismatch between functional and structural investigations. Some functional studies [6] [7] [8] [18] support the DMN definition of PCC (PGM and A19M areas) and PPC (LIP and MIP areas) for the common marmoset, but neuroanatomical (cytoarchitectonic) results [14] [15] do not support it as the homologue of the human DMN.

In this study, we carefully restarted from a neuroanatomical point of view and identified two ICA components (Comp-A and Comp-B) as candidates for the DMN. Component-A (Comp-A) is the (earlier described) network that in the literature has been called either the DMN, or FPN in the common marmoset. Comp-A peaks at Paxinos’s LIP and MIP areas (of the PPC), and PGM and A19M areas (of the PCC) (Table 1, Fig. 1a). Another one, Component-B (Comp-B), has previously been called the somatomotor network (SMN) in the common marmoset [6][8]. It peaks at the PE area (of the PPC), and A23b and A31 areas (of the PCC) (Table 1, Fig. 1c). We next reviewed Liu et al.’s visual-task fMRI experiment and noticed that their marmosets were trained to reduce their saccades. The visual-task fMRI experiment may affect task-induced deactivation around the LIP and MIP areas so we performed GLM analysis with a more appropriate auditory-task fMRI [22] [23] dataset to check for deactivated regions in the marmoset cortex. We confirmed the anatomical connectivity (from retrograde tracing) between the medial prefrontal cortex (mPFC) region (A10 area) and PCC region (A23 and A31 area) and evaluated their functional connectivity through multiseed-based connectivity analysis. Here, we confirmed that the marmoset mPFC and PCC regions were not functionally connected. Through these analysis results we propose that Comp-A is the FPN and Comp-B is the DMN of the common marmoset. Finally, we performed fingerprint analysis (following Hori et al. [8]) by using several sub-cortical regions. We made comparisons of marmoset fMRI not only with human resting-state fMRI network components, but also with human task-fMRI (working memory-task and motor-task) network components. Surprisingly, we found that both Comp-A (FPN) and Comp-B (DMN) were closer to the human task-fMRI components than the human rs-fMRI components. Reciprocally, a suppressed DMN and activated FPN in the tasking human was closer to the resting common marmoset. This suggests that the marmoset may not be resting like humans do during fMRI experiments, or, based on the combination of this result and multiseed-based connectivity analysis between mPFC and PCC regions, the resting-state DMN may be underdeveloped in the common marmoset brain.

## Results

### Anatomy-based comparison of DMN regions

Fig. 1 shows the results of our anatomy-based comparison. The human DMN component (Fig. 1b, e) is visualized in between two awake marmoset ICA components (Comp-A and Comp-B) (Fig. 1a, d and c, f). To acquire the human DMN component, 200 sessions of HCP rs-fMRI data [24] were pre-processed by the CONN toolbox [25], and group ICA (MELODIC [11]) was applied to acquire 15 components from the human rs-fMRI data. The DMN component was then manually selected. For the awake marmoset ICA components, rs-fMRI data were acquired as part of the Brain/MINDS project [26] [27] and pre-processed by Statistical Parametric Mapping (SPM12) [28]. Then, 30 components were acquired by group ICA in the same manner as for the human components. The PCC region of the human DMN component peaks around Brodmann’s [24] A23 and A31 areas (Fig. 1b), however, Comp-A, which was previously called the marmoset DMN or FPN, peaks at Paxinos’s [29] [30] PGM and A19M areas on the marmoset cortex (Fig. 1a). In neuroanatomical terms, these are inconsistent results. The PPC is also inconsistent: Comp-A peaks at Paxinos’s LIP and MIP areas, but a previous study in the macaque monkey showed that the LIP receives input from many visual areas [31] and has direct neural connections to the frontal eye field (FEF) and the superior colliculus (SC), which are the centre of the saccade oculomotor system [32] [33]. Fig. 1d showed strong positive Z-score in SC area (top right), but the human case did not (Fig. 1e top right). The MIP of the macaque monkey also seems to closely resemble the function of the human medial intraparietal cortex [34]. This is why Comp-A has been repeatedly called the FPN. The PPC of the human DMN component peaks around Brodmann’s A39 area (Fig. 1b), which corresponds to the vicinity of the PG and PFG areas [35] [36] of the marmoset cortex. We systematically examined the different components generated by ICA and found what we call Comp-B to have neuroanatomically (cytoarchitectonic) better fitting regions with the human DMN. Comp-B has a peak around the A23b, A31 areas for PCC, and includes the PG, PFG areas rather than MIP, LIP for PPC (Fig. 1c). Comp-B peaks at the PE area, which would correspond to Brodmann’s A7 area of the human cortex, which is dorsal to the human A39 area, and also includes parts of the A1, A2, and A3b areas, which are related to somatosensory function. Although Comp-B did not have positive Z-score areas in the temporal lobe and mPFC regions, which are positive in the human DMN component, Comp-A also did not have peaks around them. Comp-B also shows overlapped areas in the PCC and PPC of the DMN regions based on architectonic analysis, therefore it could be a fascinating DMN candidate. In a previous study, this component was called the (dorsal medial) somatomotor network (SMN) [6] [14]. However, it peaks around the PE and A23b, A31 areas, but not A4ab (primary motor and somatosensory areas [37]); we therefore think this component does not match with the SMN.

### Task-induced deactivation of DMN regions

Task-induced deactivation was originally observed in positron emission tomography (PET) blood flow studies [13]. In early studies task-induced decreases in blood flow were largely ignored [13]. However, Shulman et al. [38] showed that task-induced decreases in blood flow were a common phenomenon in PET activation studies. Later, this phenomenon was termed the “default mode” of brain function by Raichle et al. [39]. Thus, task-induced deactivation is one of the important techniques to help identify DMN regions. Liu et al. [7] collected visual task-fMRI of the common marmoset and found that task-induced deactivation occurs around the PCC (PGM and A19M) and PPC (LIP and MIP) regions. However, we found that marmosets were trained to reduce their saccades and as a result the eye-tracking signal was reduced (see Fig. 1 of their article). This may cause deactivation around the LIP area and may give confounding results. Additionally, Gilbert et al. [40] performed a social task-fMRI experiment with two marmosets in a whole-body human-spec 3T MRI and their marmosets were not trained to reduce saccades. Results showed activations around the PCC (PGM and A19M) and PPC (LIP and MIP) regions for both the face-to-face and movie watching paradigms. This is inconsistent with Liu et al.’s result.

Therefore, we propose that an auditory-based investigation of task-induced deactivation, rather than visual-based, may be more appropriate. For the human case, a passive sentence listening task showed significant deactivation in the PCC and mPFC regions [22], and a tone discrimination task showed significant deactivation in the PCC, PPC and mPFC regions [23]. Recently, Gilbert et al. performed an auditory task-fMRI experiment (a passive marmoset vocalization stimuli) with the common marmoset [41], but they did not visualize a surface mapping of the task-induced deactivation across the brain. We acquired their auditory task-fMRI data and performed GLM analysis to investigate the details of the task-induced deactivation (Fig. 2). The human and common marmoset have different peak times in their hemodynamic response functions (HRF); 5-6 seconds for the human [42] and around 3.1 seconds for the marmoset [43]. The canonical HRF used for GLM analysis was characterized by two gamma functions. Fig. 2a shows example canonical HRFs for the marmoset and the human. The design matrix for GLM analysis is simple, with one variable, four nuisance variables and an intercept (Fig. 2b). Auditory stimuli minus resting contrast (auditory stimuli > rest) were used for the analysis. We could successfully reproduce the activated auditory related regions of Gilbert et al.’s result, such as the inferior colliculus, medial geniculate nucleus, and auditory cortex (Fig. 2c). Fig. 2d shows a surface mapping of the GLM analysis for the auditory task-fMRI, and the auditory cortex showed task-induced activation (white arrow). From this confirmation we further investigated the task-induced deactivation. The VIP, LIP, and A19M areas did not show peak deactivation during the auditory-based task (Supplementary Table 1 gives detailed voxel rates). Instead of these areas, PEC and PE (red arrow), A23b and A31 (yellow arrow), PG, PFG (cyan arrow) and part of MIP, V2 were deactivated by the task. This result is roughly consistent with task-induced deactivation in macaque monkeys [5] in A23, A31, PEa and PGm except A24/32, A23v, A9/46d and A8b. Comp-A has a positive group ICA result in A23V, A19M, PGM, MIP, VIP, LIP, OPt, PG and part of A8b, while Comp-B is positive in A23, A31, PE, PF, PFG, A1/2, A3b and part of A8b. Although both components showed several overlapping areas between component and task-induced deactivation, A23, A31 and PE appear as consistent areas between common marmoset and macaque monkeys, therefore we prefer Comp-B as a more suitable DMN component.

**Fig. 2.**
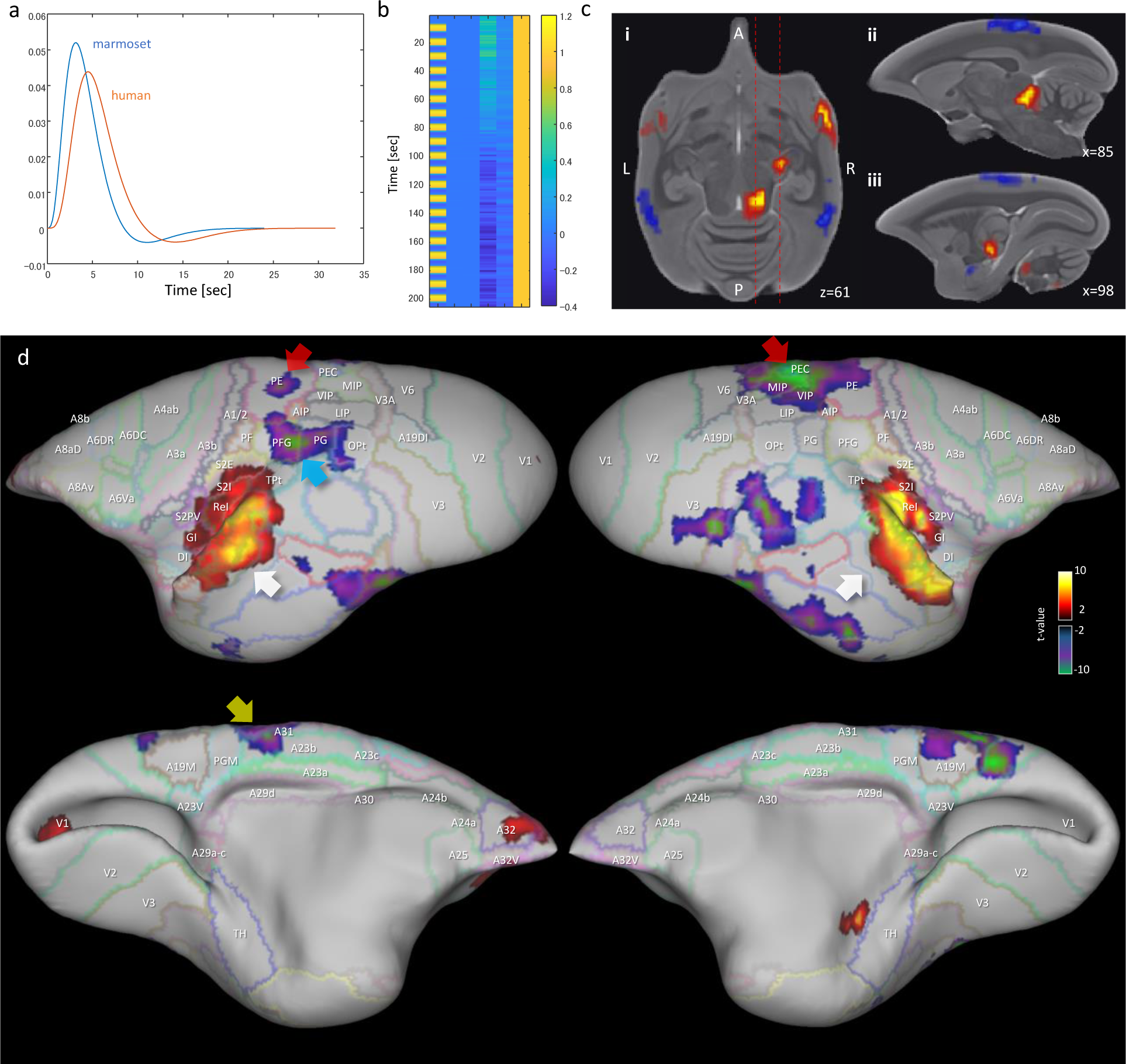
GLM analysis results of awake marmoset passive auditory task-fMRI. **a**, Canonical hemodynamic response function (HRF) for the marmoset and human. **b**, Example design matrix for GLM analysis (TR=3 seconds). **c**, GLM analysis result (auditory stimuli > rest) of sub-cortical regions. i) Horizontal plane (z=61) of marmoset brain shows several activated regions. ii) Sagittal plane (x=85) shows activated region of inferior colliculus. iii) Sagittal plane (x=98) shows activated region of medial geniculate nucleus. **d**, GLM analysis result (auditory stimuli > rest) mapped to the marmoset cortical surface. White arrow shows activated region of auditory cortex. Red arrow shows deactivated region of PEC, PE. Yellow arrow shows deactivated regions of A23b, A31. Cyan arrow shows the deactivated region of PFG.

### The functional connectivity of mPFC does not match the anatomical connectivity

The functional connectivity of the mPFC region of the common marmoset has been investigated in works such as [7] [21]. These showed that the marmoset does not have functional and anatomical connections between the PCC and PPC regions. Since we propose a new DMN candidate, Comp-B, we also investigated the functional and anatomical connectivity between DMN regions through multiseed-based connectivity analysis. Fig. 3a shows analysis results for human and awake marmoset resting-state fMRI. The human mPFC (parts of A9, A10, and A32) seeds showed significant functional connectivity with other DMN regions (Fig. 3a top), whereas the marmoset mPFC (parts of A9, A10, and A32) did not show significant functional connectivity at all. This result is consistent with Liu et al.’s [7]. We also checked the functional connectivity of the marmoset PCC (parts of A23a, A23b, and A31) and it did not show significant functional connectivity to the mPFC. However, based on the marmoset cortex retrograde tracing results from the Marmoset Brain Connectivity Atlas [44], anatomical connections were observed between A10 and A23a (Fig. 3b). We observed weak functional connectivity (around t-value=3) from the marmoset mPFC to PCC regions, but these connections disappeared after Bonferroni correction. In the human case, we observed strong connections even after family-wise error rate correction, thus the marmoset PCC region has structural connections to mPFC, but the two are functionally separated.

**Fig. 3.**
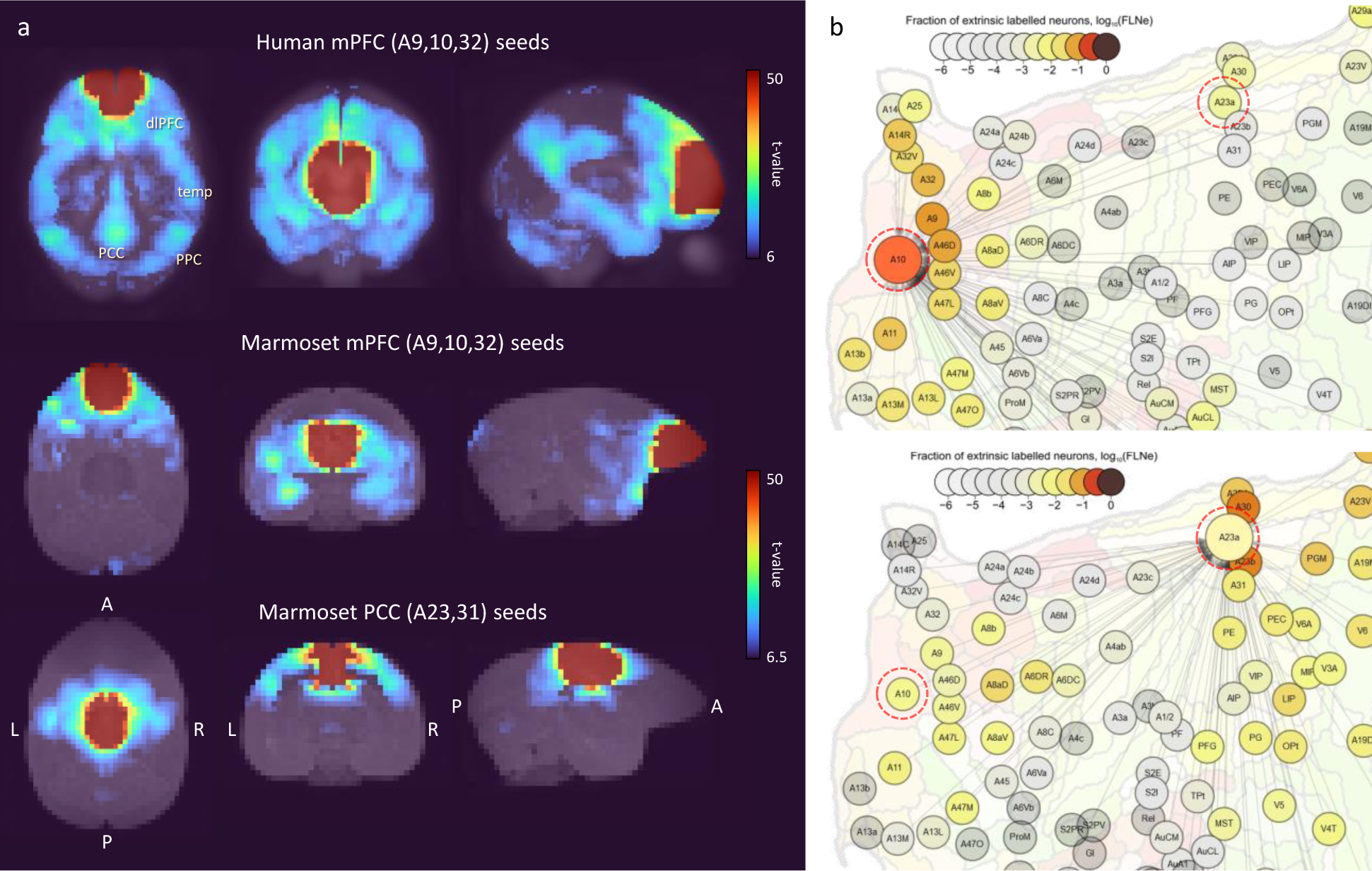
Multiseed-based connectivity analysis results of human and awake marmoset resting-state fMRI. **a**, Three-dimensional maximum projection of t-values from multiple seeds are shown. (Top) Result for seeds in human mPFC (part of A9, A10 and A32 areas). (Middle) Result for seeds in marmoset mPFC (parts of A9, A10, and A32). (Bottom) Result for seeds in marmoset PCC (parts of A23a, A23b, and A31). **b**, Retrograde tracing results of marmoset cortex from the Marmoset Brain Connectivity Atlas [44]. (Top) Injection point in area A10. (Bottom) Injection point in area A23a.

### Fingerprint analysis of resting and task state FPN and DMN

Hori et al. [8] applied fingerprint analysis [17] to analyze the correspondence between marmoset and human ICA components. To apply fingerprint analysis, they used 14 sub-cortical fingerprints (right hemisphere regions): Caudate (CAU); putamen (PUT); hippocampus (HIPPO); amygdala (AMY); superior colliculus (SC); inferior colliculus (IC); and a set of thalamic ROIs (regions of interest), namely, the lateral geniculate nucleus (LGN), anterior (ANT), laterodorsal (LD), mediodorsal (MD), ventral anterior (VA), ventral lateral (VL), ventral posterior (VP), and pulvinar (PUL) (Supplementary Fig. 1).

Topological features of sub-cortical regions are well preserved between the marmoset and human (Supplementary Fig. 1), and regional functions are also assumed to be homologous among primate species. We used these same fingerprints where the sub-cortical ROIs were taken from the Brain/MINDS 3D Marmoset Reference Brain Atlas 2019 [45] for the marmoset, and the ALLEN HUMAN REFERENCE ATLAS – 3D, 2020 [46] for the human. The correlation between component time-series (resting marmoset FPN/DMN and resting/task human FPN/DMN) and voxel time-series in sub-cortical regions was calculated for all marmoset and human sessions. A mixed-effects model was applied for group analysis and the t-value of each voxel was calculated by one-sample t-test [47]. Fig. 4 shows the fingerprint analysis result between awake resting marmoset and resting/task human ICA components. The 3D maximum projection of t-values in sub-cortical voxels are shown in Fig. 4a and b, with the top row showing the component-to-voxel correlation results of resting marmoset FPN/DMN components. The middle row shows the resting human FPN/DMN components, where we can see a component-to-voxel correlation difference between resting marmoset and resting human. It is known that the hippocampus is involved in the human DMN (Fig. 1e and [1]), and Comp-A showed a slightly stronger correlation in the hippocampus (Fig. 4a, yellow arrow), this was also shown in the resting human DMN component (Fig. 4b, yellow arrow). If we consider only the resting human components, Comp-A fits better to the human DMN and Comp-B fits better to the human FPN. However, if we consider the task human components, Comp-A fits better to the human working memory-task FPN and Comp-B fits better to the human working memory-task DMN. This opposing result is very confusing. However, when we used the mean t-values to quantify the fingerprints of the 14 sub-cortical ROIs, Fig. 4c and d show a clear difference between the marmoset and human resting FPN/DMN components, and the human task components were closer to the resting marmoset components. Also, considering neuroanatomical (cytoarchitectonic) evidence, we think that Comp-B is more suitable for the DMN. The resting human FPN component was less active and uncorrelated with the SC, while the resting marmoset FPN was more active and correlated with the SC, and the marmoset was most likely alert to its surroundings. Our results suggest that marmosets may not be ‘resting’ like a human does during an fMRI experiment, or the resting-state DMN may be underdeveloped in the common marmoset brain.

**Fig. 4.**
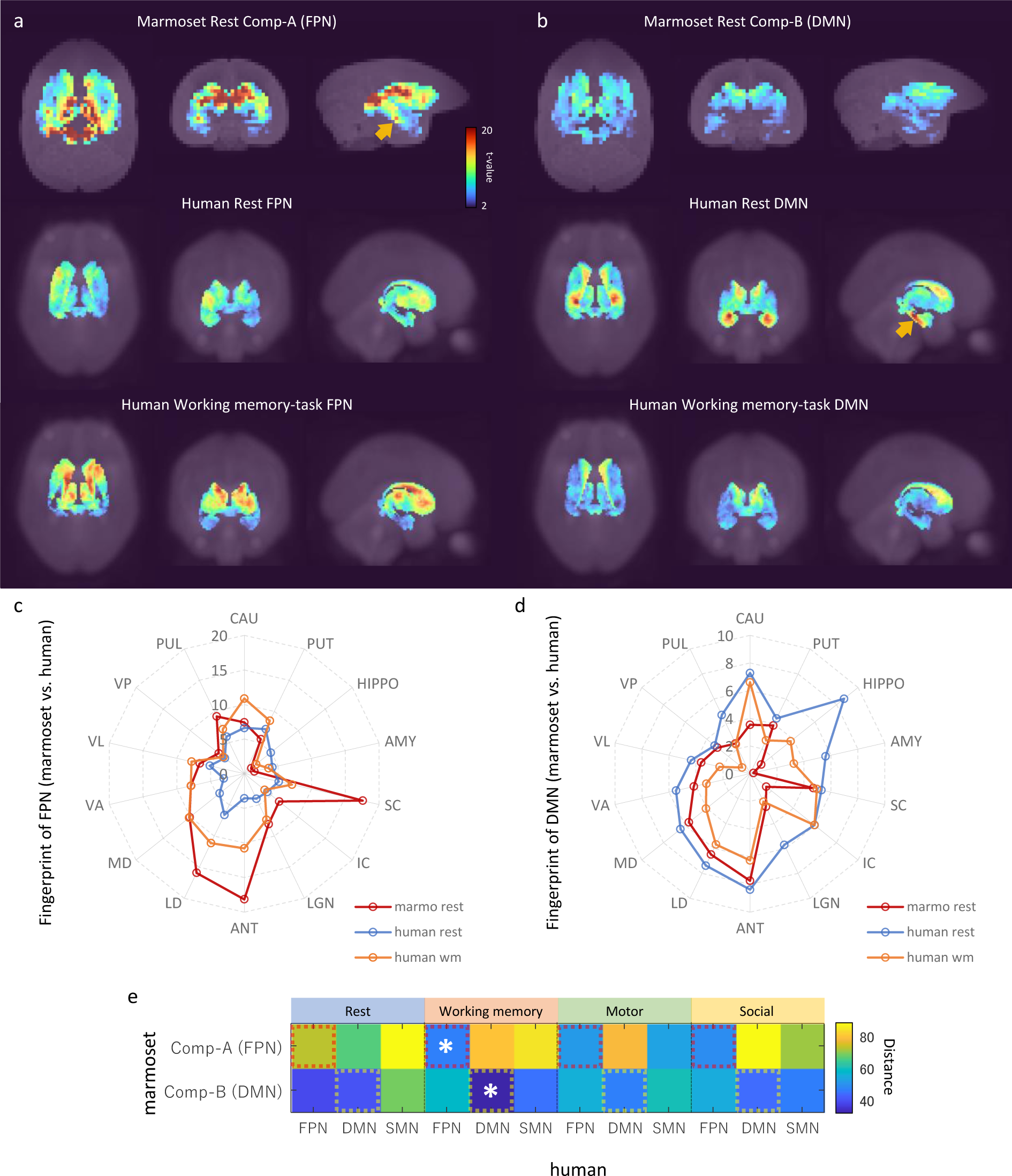
Fingerprint analysis result between awake resting marmoset and resting/tasking human ICA components. **a**, Example fingerprint result of three-dimensional maximum projection of t-values. Correlation between FPN component time-series and voxel time-series in sub-cortical regions was calculated. The closest two ICA components are shown at the top row (awake resting marmoset) and bottom row (working memory-task human). The middle row shows resting human for reference. t-value color bar is the same for all. **b**, Example fingerprint result of three-dimensional maximum projection of t-values (DMN). The bottom row shows working memory-task human. **c**, Radar chart of FPN fingerprint result of 14 sub-cortex regions (resting marmoset, resting human, and working memory-task human). **d**, Radar chart of DMN fingerprint result of 14 sub-cortex regions (resting marmoset, resting human, and working memory-task human). **e**, Fingerprint distance results between awake resting marmoset components and resting/task human components. White asterisks show the closest components from Comp-A and Comp-B.

The Manhattan distance [8] between resting marmoset FPN/DMN and resting/task human FPN/DMN components was calculated using the fingerprints of the 14 sub-cortical ROIs (Fig. 4e, and the extra marmoset ICA component version is available in Supplementary Fig. 3). Hori et al. [8] showed that marmoset Comp-A was closer to the human DMN component than to the FPN or SMN components in the resting-state and our results have been consistent with theirs. However, Comp-A was closer to the human FPN components than to the human DMN or SMN components in all three task states (working memory, motor, and social tasks). Furthermore, our proposed resting marmoset DMN component, Comp-B, was consistently closer to the human DMN components than to the human FPN or SMN components in the three task states. Our fingerprint analysis also indicated that the marmoset’s Comp-B (named dorsal medial SMN in a previous study [6] [14]), is not close to the resting/task human SMN components.

Finally, we investigated the human working memory task-fMRI data when compared with marmoset resting-state fMRI data (Fig. 5). Within each run of the working memory task, 4 different stimulus types were presented in separate blocks. Also, within each run half of the blocks used a 2-back working memory task and half used a 0-back working memory task (as a working memory comparison) [24]. The GLM result of 2-back versus 0-back contrast was then mapped onto a 3D digital brain surface (Fig. 5a). This task deactivated DMN regions, such as PCC (A23, red arrow), PPC (A39, white arrow) and mPFC (A9 and A10, yellow arrow). Even under this condition, we were able to observe the FPN and DMN ICA components (Fig. 5b and c). However, compared with the human resting-state DMN component (Fig. 1b), the activity in PCC (A23) and the inferior part of PPC (A39) became less apparent, probably due to task-induced deactivation. Based on our fingerprint analysis, the resting marmoset DMN component Comp-B was the closest to this human DMN component (Fig. 5c) over other resting/task human components. Thus, the sub-cortical activity of the resting marmoset DMN may be closer to that of the deactivated human DMN, or that a human-like activated DMN may not really be the default mode for the marmoset.

**Fig. 5.**
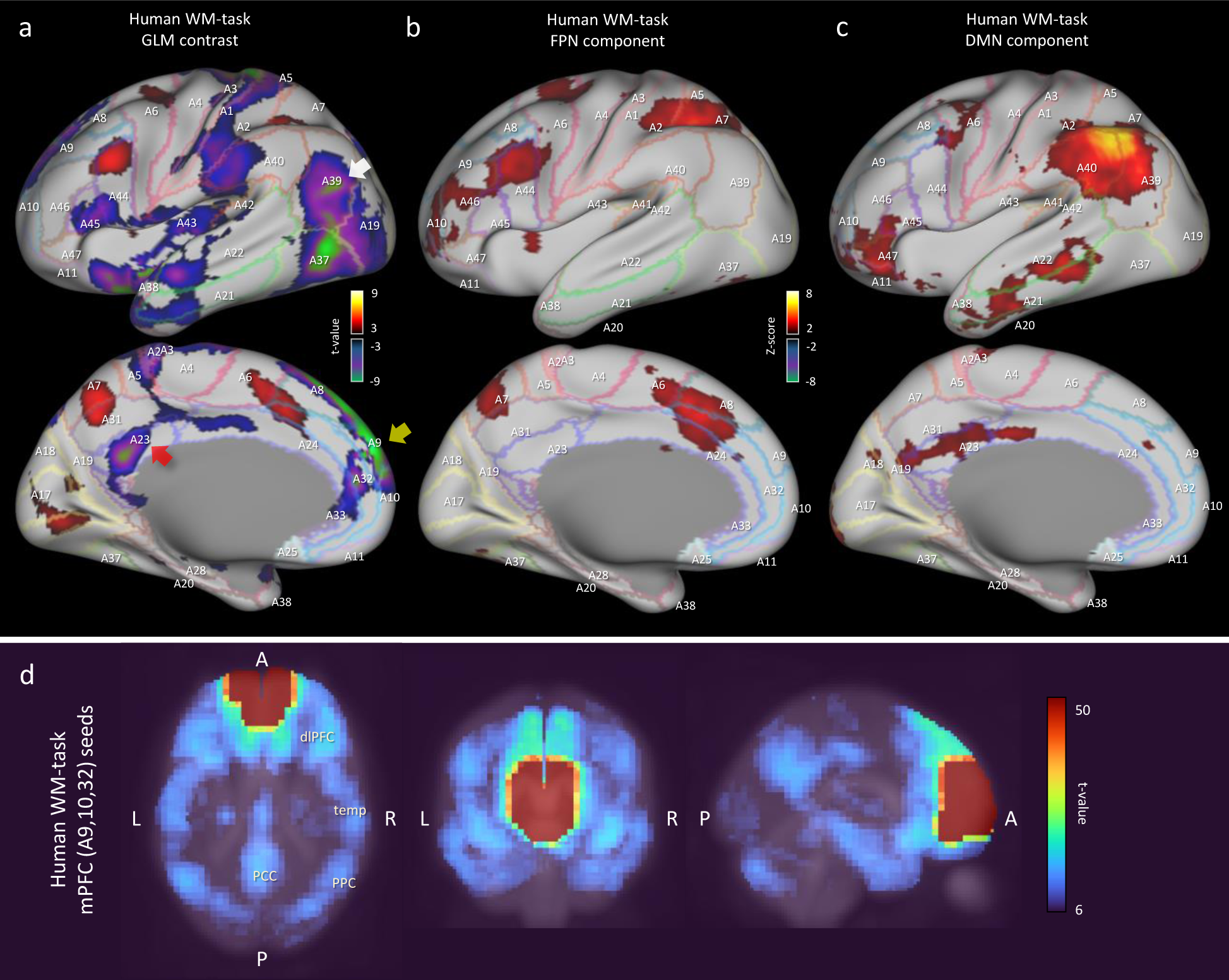
Analysis results of human working memory-task fMRI data. **a**, Left cortical surface of the human brain. GLM result of 2-back versus 0-back contrast mapped onto the surface. The t-value range is 5 to 9.1 for positive, -5 to -10.6 for negative. **b**, Left cortical surface of the human brain. Human working memory-task FPN component is mapped onto the surface. Z-value range is 3 to 15 for positive, -3 to -15 for negative. **c**, Left cortical surface of the human brain. Human working memory-task DMN component is mapped onto the surface. **d**, Multiseed-based connectivity analysis result of human working memory-task fMRI using mPFC seeds (part of A9, A10 and A32 areas). Three-dimensional maximum projection of t-values from multiple seeds are shown.

## Discussion

The DMN has a characteristic shape (Supplementary Fig. 2), such as well separated PCC, PPC and dlPFC regions, but defining it in the marmoset remains a challenge. Based on anatomy, Brodmann’s areas vary widely across primate species. In particular, the ratio of the size of the visual cortex to the total cortex varies greatly [48] [49], and the remaining sensory, motor, and functional cortex are generally more to the anterior side in the common marmoset. For this reason, the A23 and A31 cortical areas (except A23V) are much more anterior than in the human. Although the PCC and PPC regions are located posterior in the human, these regions of the marmoset are not always in a posterior location (see Fig. 1, which shows the human and marmoset A23 and A31 locations). As a result of neuroanatomical verification (Fig. 1), auditory based task-induced deactivation (Fig. 2), and fingerprint analysis by sub-cortical regions (Fig. 4), we determined that Comp-A is the FPN and Comp-B is the DMN in the common marmoset brain. Comp-A has been mentioned in various papers with a debate over it being the DMN or the FPN [6] [14] [7] [8] [18] [20] [21] and we see several pieces of evidence that suggest it might be the DMN, such as part of MIP showing task-induced deactivation (Fig. 2d), A23V showing task-induced deactivation in macaque monkeys [5], and fingerprint analysis between the resting human DMN and resting marmoset Comp-A being closer than with Comp-B. Although we propose that Comp-B is more suitable for the DMN component through several lines of evidence, further investigation will be required to definitively determine the DMN in the common marmoset brain.

Our results suggest that the structure of large-scale brain network components of the human, such as the DMN, should not always be relied upon for defining the equivalents in species such as the common marmoset. The human brain is much larger than that of the marmoset (190-fold difference in weight [50]) and is gyrified with deep sulci. Due to this the BOLD (blood oxygenation level-dependent) signal is well separated between regions. For example, supplementary Fig. 4 shows a human SMN component with a clear boundary around the somatomotor (A4) and somatosensory cortex (A1/2, 3). However, the marmoset cortex is smooth and relatively small, and Comp-B showed ambiguity in regional boundaries around PE, A1/2, A3, and A4ab, even in ultrahigh field (9.4T) fMRI data. The temporal resolution of fMRI scans of the marmoset brain is low compared with the human. Current marmoset data has TR=2.0 seconds and group ICA was obtained from 140 frames × 48 sessions. This small dataset may result in insufficient component decomposition. A higher temporal resolution (or larger frame number) and larger session data would be required to correct this ambiguity for the marmoset.

Garin et al. [21] also showed the differences between the human DMN and the non-human primate FPN (Comp-A) in resting state fMRI using fingerprint analysis. They showed that the human PCC highly correlated with the PPC and the mPFC, but the non-human primate PCC is highly correlated with the PPC and the dlPFC. Furthermore, the human mPFC highly correlated with PPC, Temp, and PCC, but the non-human primate PCC does not. However, marmoset FPN component was originally named by Ghahremani et al. [14] based on seed analysis of superior colliculus, and their claim was supplanted by more direct evidence from visual-fixation task-induced deactivation of the DMN [7]. Thus, Garin et al. could not exclude either the front temporal network (FTN) or the FPN as homologous candidates to the human DMN. We directory challenged this issue, and based on auditory task-induced deactivation we found Comp-B is more suitable as the DMN, and based on fingerprint analysis, Comp-A is closer to the (task-induced activated) human FPN. In our study, we could not find a clear FTN component by group ICA, and multiseed-based connectivity analysis of the marmoset mPFC region did not show strong connectivity with the temporal lobe (in comparison to the human case) (**Fig. 3**). Therefore, no judgment can be made regarding this network and further research is required.

The function of the DMN in the resting marmoset also resulted in questionable results as to whether it is homologous to the resting human DMN. Although marmosets might not be resting in the same manner as a human during fMRI experiments, marmosets are usually trained to become familiar with the MRI machine, and in our experiment four marmosets went through as many as 12 sessions. This training to be familiar allows us to say that the marmoset can be considered to be in a state of rest. However, we found that the resting marmoset DMN component (Comp-B) was not close to the resting human DMN component; based on fingerprint analysis we found that it was closer to the working memory-task human DMN component. The human DMN was highly suppressed in this task (Fig. 5a) and fingerprint analysis showed a low correlation in activity between the DMN component and sub-cortical voxels (Fig. 4d).

Conversely, activity in the resting human DMN correlated highly with sub-cortical voxels, and marmosets appear to have correlations somewhere in-between. Thus, the DMN of the marmoset (Comp-B) is not as active as the human in the resting state, and it implies that we should consider that a marmoset does not reach a state that could be considered the default mode for the human. The human DMN appears to have an advanced function [13] that may be underdeveloped in non-human primates (such as the common marmoset) [21]. From now on, we need to consider that the human default mode network is not the same as the default mode for the common marmoset, and possibly for other non-human primates [21] and rodent species.

## Methods

### Preprocessing of marmoset resting-state fMRI data

Awake resting-state fMRI data of the common marmoset (*Callithrix jacchus*) were acquired as part of the Brain/MINDS project [26] [27]. A Bruker BioSpec 9.4T MRI machine (Biospin GmbH, Ettlingen, Germany) was used. The experimental settings of the gradient recalled echo planar imaging (EPI) sequence were as follows: flip angle = 65, repetition time (TR) = 2,000 ms, echo time (TE) = 16 ms, pixel size = 0.7 × 0.7 mm, slice thickness = 0.7 mm, matrix size = 60 × 42 × 52, and frame length = 150.

For our experiments, T1WI, T2WI, and rs-fMRI NIfTI files of awake marmosets (3 to 6 years, 3 males and 1 female, 12 sessions per subject) (N=48) were used. Preprocessing and image registration were performed using Statistical Parametric Mapping (SPM12) [28]. Realignment was applied for NIfTI images to compensate for head movement by a least squares approach and a 6 parameter (rigid body) spatial transform. Slice timing correction was performed to correct for signal acquisition timing discrepancies in each section, and images were co-registered to the Marmoset MRI Standard Brain [51]. We removed the first 10 frames of the rs-fMRI data, and the remaining data were smoothed using a full width at half maximum (FWHM) of 1.4 mm (2 voxels) for group ICA (for compatibility with [7]). A FWHM of 2.4 mm (3.4 voxels) was used for multiseed-based connectivity and fingerprint analysis. Global mean and aCompCor [52] were applied for nuisance factor removal and a high-pass filter (1/128Hz) was applied for subsequent analyses.

### Independent component analysis of marmoset resting-state fMRI data

After preprocessing, independent component analysis (ICA) was applied to the marmoset rs-fMRI data to acquire 30 components. The number of components was chosen for compatibility with [7]. MELODIC [11] was used to obtain group ICA from 48 sessions (140 frames). Here, multi-session temporal concatenation was performed and a spatial map was obtained. Finally, the two components that were used in our study, Comp-A and Comp-B, were manually selected from the 30 components.

For surface mappings of marmoset data, we first converted NIfTI images from the Marmoset MRI Standard Brain space [51] to the Marmoset Brain Mapping V3 space [30]. Then, ‘wb_command-volume-to-surface-mapping’, included in the Connectome Workbench visualization software [53], was used to map NIfTI image data onto the marmoset cortical surface. Finally, the cortical surface (in gray), the functional data mapped to the surface, and the Paxinos label map [29] were overlaid to produce our figures.

### Multiseed-based connectivity analysis of marmoset resting-state fMRI data

Multiseed-based connectivity analysis was done by calculating the correlation coefficients between seed voxels and all other voxels. MATLAB scripts for this analysis were developed in-house and worked together with the VARDNN toolbox [54]. The seed voxels of the marmoset mPFC and PCC regions were manually edited in ITK-SNAP [55]. After calculating the correlation coefficients in each voxel from individual sessions, a mixed-effects model was applied to acquire final group results. A one-sample t-test in each voxel was performed for 2nd-level (group) analysis [47]. Bonferroni correction was then applied to correct for the familywise error (FWE) rate and *t*-value threshold (*t* >6.48 in Fig. 3) were applied to acquire significantly correlated voxels.

### Fingerprint analysis of marmoset resting-state fMRI data

Fingerprint analysis [17] was used to analyze the correspondence between marmoset and human ICA components. To apply fingerprint analysis, we used 14 sub-cortical regions as fingerprints (Supplementary Fig. 1) from the Brain/MINDS 3D Marmoset Reference Brain Atlas 2019 [45]. The correlation between component time-series (resting marmoset FPN/DMN) and voxel time-series in sub-cortical regions was calculated for all marmoset sessions. A mixed-effects model was applied for group analysis and the t-value of each voxel was calculated by one-sample t-test. Mean t-values were used to quantify the fingerprints of the 14 sub-cortical ROIs. Finally, the Manhattan distance between resting marmoset FPN/DMN and resting/task human FPN/DMN components was calculated using all 14 fingerprints.

### Preprocessing of marmoset auditory task-fMRI data

Gilbert et al. [41] performed an auditory task-fMRI experiment with the common marmoset. We used their auditory task-fMRI data to investigate task-induced deactivation. Three functional time courses were acquired from two awake marmosets (named M3 and M4). Details of the data are orientation: axial, resolution: 500-μm isotropic, FOV: 48 × 48 mm, number of slices: 42, number of volumes: 205, TE: 15 ms, BW: 400 kHz, flip angle: 40◦, acceleration rate: 2 (left-right).

T2WI and task-fMRI NIfTI files (N=6) were used for registration. Preprocessing and registration were performed using Statistical Parametric Mapping (SPM12) [28]. SPM12 registered NIfTI images to the Marmoset MRI Standard Brain [51], and task-fMRI data was smoothed using a FWHM of 1.7 mm (3.4 voxels). The preprocessed task-fMRI data was then used for GLM analysis.

### GLM analysis of marmoset auditory task-fMRI data

GLM analysis was used to investigate the details of the task-induced deactivation. The canonical haemodynamic response function (HRF) used for GLM analysis was characterized by two gamma functions with peak time around 3.1 seconds (for the marmoset) [43]. A simple GLM design matrix was used with one variable, four nuisance variables and an intercept. Data for the first variable were created by convolution of the canonical HRF from block car designs corresponding to sound stimuli. Data for the four nuisance variables were calculated from the average values for each time point of the white matter, CSF, all brain voxels, and the average signal over all voxels. A high-pass filter (1/128Hz) was applied to the target variable and first variable, then a Tukey taper (taper size = 8) was used for GLM pre-whitening [56]. The mixed-effects model was used for group analysis and the t-value of each voxel was calculated by 2^nd^-level analysis of OLS regression with a Tukey taper. Then, we applied a voxel-wise primary threshold (uncorrected p-value < 0.001 and *t*>4.14) to obtain significantly activated or deactivated voxels [57], and a cluster-extent threshold (k>69 voxels and FWE corrected p-value < 0.049) was applied to acquire significant clusters under multiple comparisons.

### Preprocessing of HCP resting-state fMRI data

Resting-state fMRI data from the WU-Minn HCP consortium (the S500 release [24]) were used for our experiments. Scanning used a customized SC72 gradient insert and a body transmitter coil with 56 cm bore size, and data was saved in NIfTI format. Experimental settings of the gradient-echo echo-planar imaging (EPI) sequence were as follows: flip angle = 52, repetition time (TR) = 720 ms, echo time (TE) = 33.1 ms, pixel size = 2 × 2 mm, slice thickness = 2 mm, matrix size = 104 × 104 × 90, multiband factor = 8, and frame length = 1,200. More information on the resting-state parameters can be found at the HCP website: (https://www.humanconnectome.org/storage/app/media/documentation/s500/HCP_S500_Release_Reference_Manual.pdf).

T1WI, T2WI, and rs-fMRI NIfTI files from the S500 release were downloaded and a total of 200 sessions (50 male subjects × 2 sessions, 50 female subjects × 2 sessions) were used in our experiments. The CONN toolbox [25] was used for preprocessing. CONN performed the realignment and co-registration of NIfTI images to the standard Montreal Neurological Institute (MNI) brain space. The first 10 frames of the rs-fMRI data were removed and the remaining data were smoothed using a FWHM of 4 mm (2 voxels) for group ICA (for compatibility with [7]), and a FWHM of 6.8 mm (3.4 voxels for compatibility with the marmoset data) for multiseed-based connectivity and fingerprint analysis. Global mean and aCompCor [52] were applied for nuisance factor removal and a high-pass filter (1/128Hz) was then applied for subsequent analyses.

### Preprocessing of HCP task-fMRI data

Three types of task-fMRI data (working memory, motor, social) were obtained from the WU-Minn HCP consortium (the S500 release [24]). We chose these data because the motor task is a very basic task for fMRI studies, the social task was used in previous studies for the marmoset [7] [40], and we assumed the working memory task deactivates the DMN. The experimental settings of the EPI sequence were the same as for the resting-state fMRI data. T1WI, T2WI, and rs-fMRI NIfTI files from the S500 release were downloaded and a total of 200 sessions (100 male subjects, 100 female subjects) were used in our experiments. The CONN toolbox [25] was used for task-fMRI data preprocessing. CONN registered NIfTI images to the standard Montreal Neurological Institute (MNI) brain space. Data were smoothed using a FWHM of 4 mm (2 voxels) for group ICA, and a FWHM of 6.8 mm (3.4 voxels for compatibility with the marmoset data) for multiseed-based connectivity and fingerprint analysis. A high-pass filter (1/128Hz) was applied for subsequent analyses.

### Independent component analysis of HCP resting/task fMRI data

After preprocessing, group ICA was applied to acquire 15 components from the human rs-fMRI data. We systematically checked several different numbers of components - 5/10/15/20/30 - and decided that 15 components were appropriate. For example, the default mode network became separated into two components if 30 components were chosen. MELODIC [11] was used to obtain group ICA from 200 sessions. Here, multi-session temporal concatenation was performed and a spatial map was obtained. Finally, the DMN, FPN and SMN components used in our study were manually selected from the 15 resting/task fMRI data components.

For surface mappings of human data, the command ‘wb_command-volume-to-surface-mapping’ of the Connectome Workbench visualization software [53] was used to map NIfTI image data onto the human cortical surface. Finally, the cortical surface (in gray), the mapped functional data, and the Brodmann label mapping (included in the HCP data) were overlaid to produce our visualizations.

### Multiseed-based connectivity analysis of HCP resting/task fMRI data

The procedure of multiseed-based connectivity analysis of HCP resting/task fMRI data was the same as for the marmoset. A *t*-value threshold (*t* >5.96 in Fig. 3 and Fig. 5) was applied to acquire significantly correlated voxels.

### Fingerprint analysis of HCP resting/task fMRI data

Fingerprint analysis [17] was used to analyze the correspondence between marmoset and human ICA components. To apply fingerprint analysis, we used 14 sub-cortical fingerprints (Supplementary Fig. 1) from the ALLEN HUMAN REFERENCE ATLAS - 3D, 2020 [46] for the human data. The correlation between component time-series (resting or task human FPN/DMN) and voxel time-series in sub-cortical regions was calculated for all data. The procedure to acquire t-values of 14 sub-cortical ROIs was the same as for the marmoset. Finally, the Manhattan distance [8] between resting marmoset FPN/DMN and resting/task human FPN/DMN components was calculated using the fingerprints of the 14 sub-cortical ROIs.

### GLM analysis of HCP task-fMRI data

The design matrix for GLM analysis was composed of several contrast variables, four nuisance variables and an intercept. Data for the contrast variables were created by convolution of the canonical HRF with block car designs corresponding to task stimuli. Data for the four nuisance variables were calculated from the average values at each time point of the white matter, CSF, all brain voxels, and all voxels of the volume. A high-pass filter (1/128Hz) was applied to the target and contrast variables, then a Tukey taper (taper size = 8) was used for GLM pre-whitening [56]. The mixed-effects model was applied for group analysis and the t-value of each voxel was calculated by a 2^nd^-level analysis of OLS regression with a Tukey taper. We applied a voxel-wise primary threshold (uncorrected p-value < 0.001 and *t*>3.10) to obtain significantly activated or deactivated voxels [57], and a cluster-extent threshold (k>55 voxels and FWE corrected p-value < 0.049) was applied to acquire significant clusters under multiple comparisons.

### Statistical information

For multiseed-based connectivity analysis, a mixed-effects model was used for group analysis. A one-sample t-test for each voxel was performed as a 2nd-level (group) analysis. Statistical significance was set at *p*<0.05. Bonferroni correction was then applied to correct for the familywise error (FWE) rate and a *t*-value threshold was applied to acquire significantly correlated voxels.

For GLM analysis, a mixed-effects model was used for group analysis and the t-value of each voxel was calculated by 2^nd^-level analysis of OLS regression with a Tukey taper. We applied a voxel-wise primary threshold (uncorrected p-value < 0.001 and *t*>4.14) to obtain significantly activated or deactivated voxels, and a cluster-extent threshold (k>69 voxels and FWE corrected p-value < 0.049) was applied to acquire significant clusters for the HCP task-fMRI data. For the marmoset task-fMRI data, a voxel-wise primary threshold (uncorrected p-value < 0.001 and *t*>3.10) and a cluster-extent threshold (k>55 voxels and FWE corrected p-value < 0.049) were applied.

## Data Availability

The human rs-fMRI data analyzed during the current study are available from the HCP website: https://www.humanconnectome.org/.

Pre-processed auditory task fMRI data of the common marmoset is available at https://doi.org/10.5281/zenodo.7827225.

All other datasets generated and/or analyzed during the current study are available from the corresponding author on reasonable request.

## Code Availability

The code used in the current study (GLM and fingerprint analysis for the common marmoset) are provided in open source and publicly available from https://github.com/takuto-okuno-riken/oku2023dmn.

## Authors’ Contributions

T.O. conceived of the presented idea. T.O. developed the theory, performed the computations. T.O. and A.W. discussed the results and contributed to the final manuscript.

## Acknowledgements

The authors wish to thank Dr. Cirong Liu for the valuable discussions on the default mode network.

This research was supported by the program for Brain Mapping by Integrated Neurotechnologies for Disease Studies (Brain/MINDS) from the Japan Agency for Medical Research and Development, AMED. Grant number: JP15dm0207001.

Data were provided [in part] by the Human Connectome Project, WU-Minn Consortium (Principal Investigators: David Van Essen and Kamil Ugurbil; 1U54MH091657) funded by the 16 NIH Institutes and Centers that support the NIH Blueprint for Neuroscience Research; and by the McDonnell Center for Systems Neuroscience at Washington University.

## Ethics declarations

All experimental procedures were approved by the Experimental Animal Committee of RIKEN or by the Experimental Animal Committee of the National Center of Neurology and Psychiatry. The marmosets were handled by the “Guiding Principles of the Care and Use of Animals in the Field of Physiological Science” formulated by the Japanese Physiological Society.

## Supplementary Information

**Supplementary Fig. 1.**
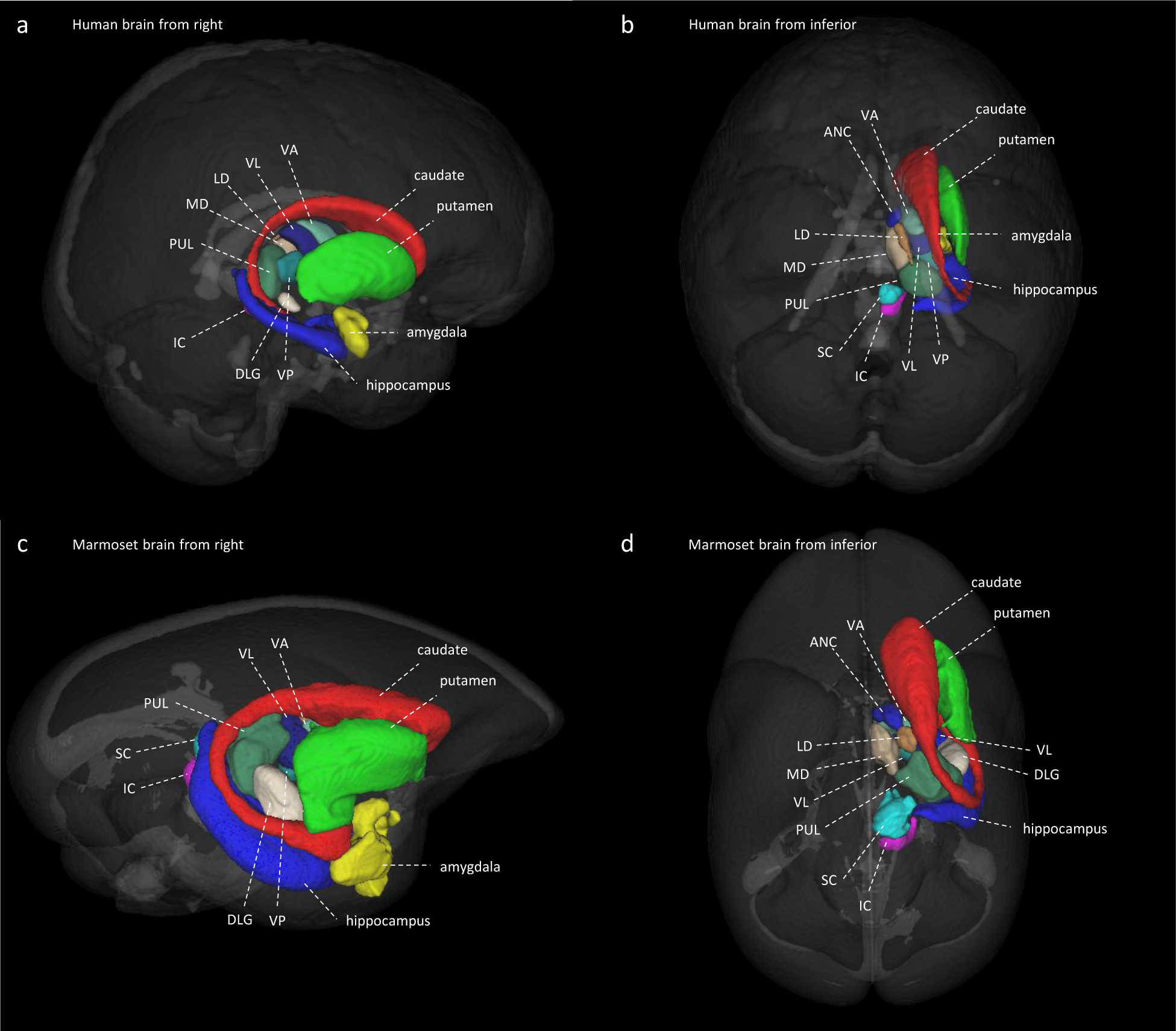
Sub-cortical ROIs of human and marmoset for fingerprint analysis. **a,** Sub-cortical ROIs of human brain from right. **b,** Sub-cortical ROIs of human brain from inferior (only right side is presented). **c,** Sub-cortical ROIs of marmoset brain from right. **d,** Sub-cortical ROIs of marmoset brain from inferior (only right side is presented).

**Supplementary Fig. 2.**
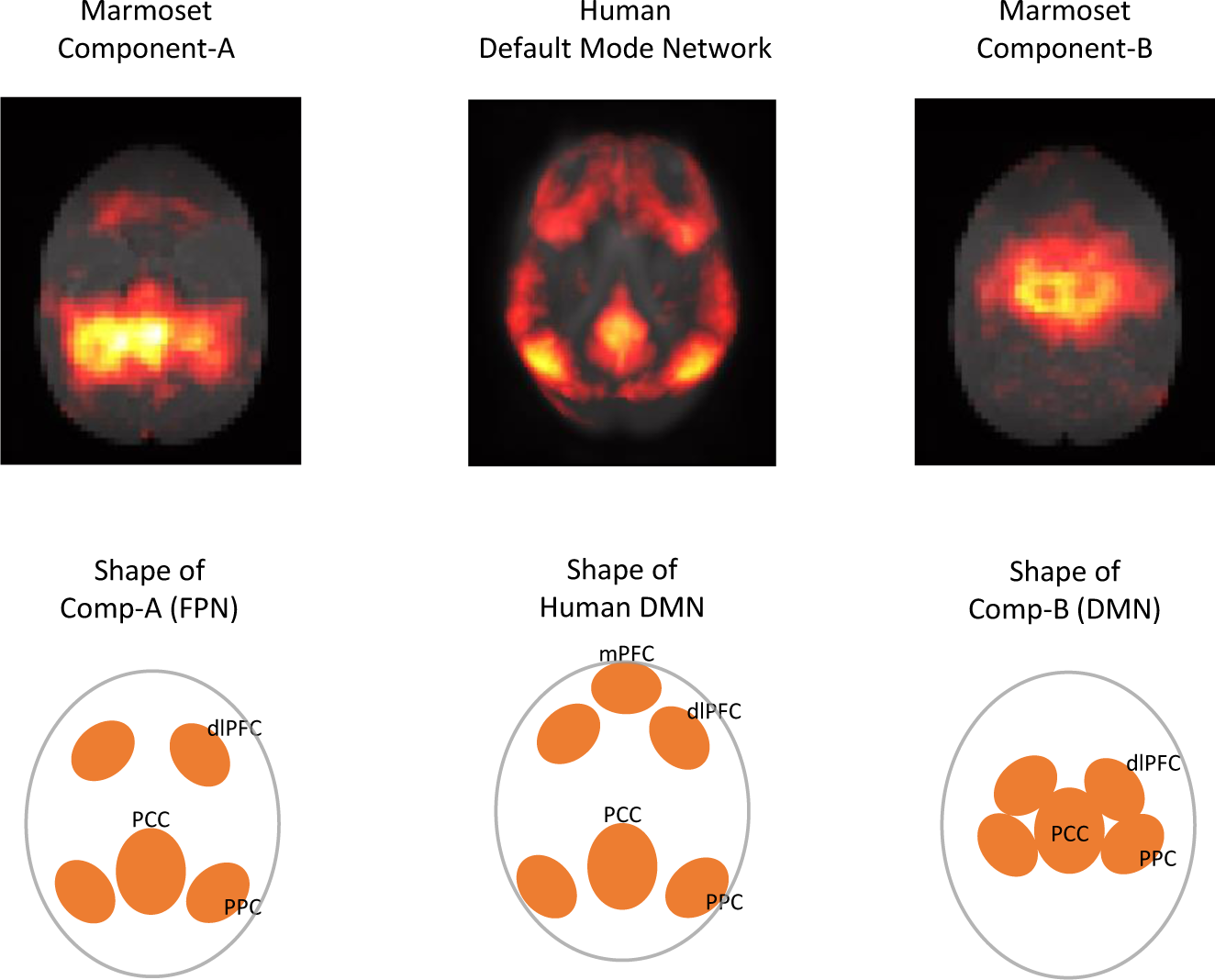
Shape of DMN component. (left) marmoset component-A and its shape in the horizontal plane. (center) human DMN component and its shape in the horizontal plane. (right) marmoset component-B and its shape in the horizontal plane.

**Supplementary Fig. 3.**
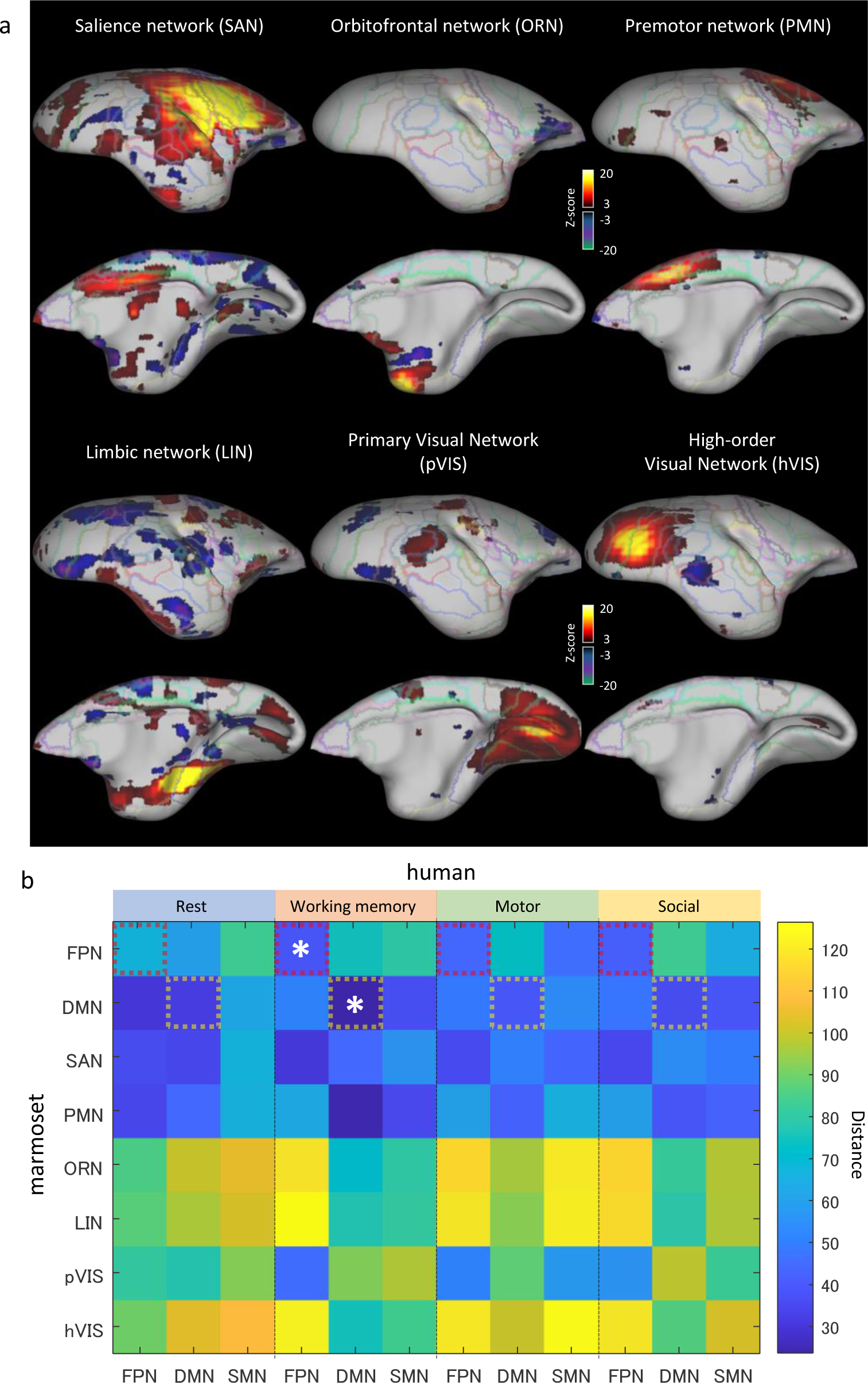
Fingerprint analysis result between awake resting marmoset and resting/tasking human ICA components. **a,** Right cortex of a marmoset surface image. (top) lateral side, (bottom) medial side. Extra components of awake marmoset ICA are mapped onto the brain surface. Z-score range is 3 to 30 for positive, -3 to -30 for negative. **b,** Fingerprint distance results between awake resting marmoset components (extra version) and resting/tasking human components. White asterisks show the closest components from Comp-A and Comp-B. Marmoset FPN, SAN and pVIS show a similar tendency. Marmoset DMN and PMN show a similar tendency.

**Supplementary Fig. 4.**
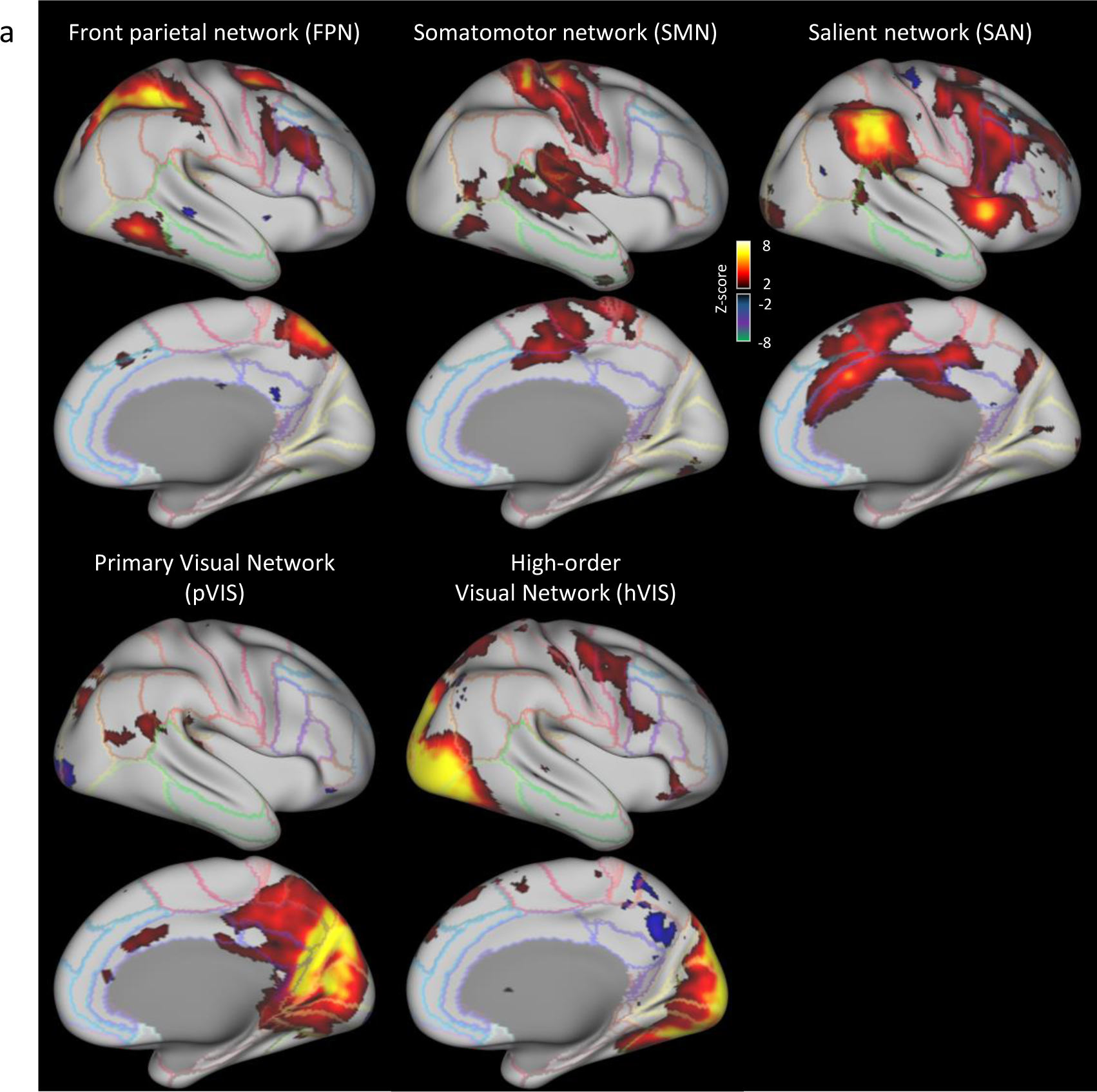
Human brain other network components. **a,** Right cortical surface of the human brain. Human resting-state network components are mapped onto the brain surface. Z-score range is 2 to 8 for positive, -2 to -8 for negative.

**Supplementary Table 1.** GLM analysis results of awake marmoset passive auditory task-fMRI.

| ROI name<br>(left<br>cortex) | significant<br>voxel rate | significant<br>voxel num | ROI<br>voxel<br>num | mean of<br>significant<br>voxels | ROI name<br>(right<br>cortex) | significant<br>voxel rate | significant<br>voxel num | ROI<br>voxel<br>num | mean of<br>significant<br>voxels |
| --- | --- | --- | --- | --- | --- | --- | --- | --- | --- |
| PE | 4.21% | 11 | 261 | -4.89 | PE | 24.72% | 66 | 267 | -4.90 |
| PEC | 16.18% | 11 | 68 | -5.41 | PEC | 57.35% | 39 | 68 | -4.88 |
| PF | 0% | 0 | 47 | NaN | PF | 0% | 0 | 45 | NaN |
| PFG | 53.33% | 32 | 60 | -4.94 | PFG | 0% | 0 | 64 | NaN |
| PG | 32.05% | 25 | 78 | -4.94 | PG | 0% | 0 | 76 | NaN |
| LIP | 2.31% | 3 | 130 | -4.55 | LIP | 2.42% | 3 | 124 | -4.91 |
| MIP | 0% | 0 | 69 | NaN | MIP | 23.68% | 18 | 76 | -5.14 |
| VIP | 0% | 0 | 27 | NaN | VIP | 23.08% | 6 | 26 | -5.58 |
| A23a | 0% | 0 | 73 | NaN | A23a | 0% | 0 | 66 | NaN |
| A23b | 7.35% | 10 | 136 | -4.72 | A23b | 0% | 0 | 137 | NaN |
| A31 | 23.08% | 12 | 52 | -5.32 | A31 | 12.28% | 7 | 57 | -5.05 |
| PGM | 0% | 0 | 83 | NaN | PGM | 2.50% | 2 | 80 | -4.43 |
| A19M | 0% | 0 | 99 | NaN | A19M | 15.69% | 16 | 102 | -5.21 |

